# DNA methylation profiling identifies TBKBP1 as potent amplifier of cytotoxic activity in CMV-specific human CD8^+^ T cells

**DOI:** 10.1101/2023.11.06.565829

**Authors:** Zheng Yu, Varun Sasidharan-Nair, Agnes Bonifacius, Fawad Khan, Thalea Buchta, Michael Beckstette, Jana Niemz, Philipp Hilgendorf, Beate Pietzsch, Philip Mausberg, Andreas Keller, Christine Falk, Dirk Busch, Melanie M. Brinkmann, Kilian Schober, Luka Cicin-Sain, Fabian Müller, Britta Eiz-Vesper, Stefan Floess, Jochen Huehn

## Abstract

Epigenetic mechanisms stabilize gene expression patterns during CD8^+^ T cell differentiation. However, although adoptive transfer of virus-specific T cells is clinically applied to reduce the risk of virus infection or reactivation in immunocompromised individuals, the DNA methylation pattern of virus-specific CD8^+^ T cells is largely unknown. Hence, we here performed whole-genome bisulfite sequencing of cytomegalovirus-specific human CD8^+^ T cells and found that they display a unique DNA methylation pattern consisting of 79 differentially methylated regions when compared to bulk memory CD8^+^ T cells. Among them was *TBKBP1*, coding for TBK-binding protein 1 that can interact with TANK-binding kinase 1 (TBK1) and mediate pro-inflammatory responses in innate immune cells downstream of intracellular virus sensing. Since TBKBP1 has not yet been reported in T cells, we aimed to unravel its role in virus-specific CD8^+^ T cells. *TBKBP1* demethylation in terminal effector CD8^+^ T cells correlated with *TBKBP1* expression and was stable upon long-term *in vitro* culture. TBKBP1 overexpression resulted in enhanced TBK1 phosphorylation upon stimulation of CD8^+^ T cells and significantly improved their virus neutralization capacity. Collectively, our data demonstrate that TBKBP1 modulates virus-specific CD8^+^ T cell responses and could be exploited as therapeutic target to improve adoptive T cell therapies.

## INTRODUCTION

Human cytomegalovirus (CMV) is one of the pathogens that shapes the host immune system through multiple infection phases, including (i) activation of innate immune responses during replication phase, (ii) nudging both adaptive and innate immune responses during persistence phase, and (iii) promoting immune inflation during latency phase (1). A significant risk of non-relapse mortality has been reported in association with CMV reactivation after allogeneic hematopoietic stem cell transplantation (HSCT) (2,3). Studies have shown that post-transplant virus-infected patients have a low incidence of antiviral T cells and a delayed generation of virus-responsive T cells, suggesting that functional antiviral T cells are essential for eliminating viral infections (4,5). During the initial period, following HSCT, patients suffering from CMV reactivation exhibit a rapid reconstitution of their CD8^+^ T cell populations due to clonal expansion of CMV-specific T cells, leading to an unfavourable CD4:CD8 ratio (6–8). Adoptive transfer of CMV-specific T cells has been shown to significantly reduce the risk of infection and reactivation of CMV in transplant recipients (9–13). However, CMV-specific CD8^+^ T [T(CMV)] cells show a high degree of heterogeneity and possess distinct immunological functions (14–16). Consequently, current adoptive transfer treatment strategies can still be improved to ensure optimal CD8^+^ T cell-mediated immunity.

Upon natural virus infection, naive antigen-specific CD8^+^ T cells are primed with viral antigen, undergo clonal expansion and differentiate into long-lived memory or terminally differentiated effector T cells (17,18). During this differentiation process, CD8^+^ T cells acquire cytolytic functions that involve the action of multiple transcription factors like T-bet and Eomesodermin (EOMES), which co-operate to induce the expression of various effector molecules, including interferon gamma (IFN-γ), perforin, and granzyme B (19,20). Effector CD8^+^ T cells respond to their targets directly through antigen-specific cytotoxic activity and secretion of multiple cytokines/effector molecules. The human CD8^+^ T cell population consists of various functionally distinct subsets that can be distinguished through the changes in surface expression of homing markers and co-stimulatory molecules including CCR7, CD62L, CD27, CD28, and CD45RA during the differentiation process (21,22). Consequently, these markers have been utilized to distinguish the phenotype of different human CD8^+^ T cell subsets, namely naive (T_N_), stem cell memory (T_SCM_), central memory (T_CM_), effector memory (T_EM_), and terminal effector memory T cells re-expressing CD45RA (T_EMRA_).

Accumulating evidence suggests that various epigenetic processes contribute to the T cell fate specification upon CD8^+^ T cell differentiation (23–25). Changes in DNA methylation patterns and histone marks were shown to play a significant role in altering the transcriptional programme of effector-related or stemness-related genes during differentiation of CD8^+^ T cell subsets (23,24,26). Genome-wide profiling studies of naive and memory CD8^+^ T cell subsets revealed a dynamic distribution of histone marks, including H3K4me2, H3K4me3, and H3K27me3 (27–29). In accordance with the assumption that DNA methylation in promoter regions is typically correlated with transcriptional inactivation (30,31), several studies have demonstrated that during CD8^+^ T cell differentiation a substantial loss of methylation marks in the promoter regions is followed by transcriptional activation of corresponding genes (32–34). Although DNA methylation studies typically emphasize on promoter regions, the methylation processes observed in other genomic regions, including CpG islands/clusters (CGI), non-coding intergenic regions, and gene bodies also affect the transcription network (31). Consequently, it is imperative to extend DNA methylation studies beyond classical promoter and transcriptional start site-specific analyses to gain a deeper understanding of the impact of epigenetic processes on T cell fate specification. Apparently, DNA methylation processes at distal regulatory domains, such as enhancers and CGIs, are often found to be negatively associated with both gene transcription and active histone modifications during cellular differentiation and phenotype commitment (35–37). However, a murine study reported that during CD8^+^ T cell differentiation in response to acute lymphocytic choriomeningitis virus (LCMV) infection the vast majority of differentially methylated regions (DMRs) were observed in promoter regions (32). These data suggest that the methylation landscape of DMRs can be a critical factor in the regulation of gene expression during CD8^+^ T cell differentiation.

The current study focuses on the epigenetic characterisation of T(CMV) cells with respect to genome-wide DNA methylation patterns. We found that T(CMV) cells display a unique DNA methylation pattern consisting of 79 DMRs when compared to bulk memory CD8^+^ T (T^mem^) cells. Among these epigenetic changes, we identified a DMR in the *TBKBP1* gene, which was further studied since its role for the cytotoxicity of CD8^+^ T cells is unknown. Intriguingly, *TBKBP1* was found to be highly expressed in T^EMRA^ cells isolated from the peripheral blood of CMV-seropositive donors, and its expression significantly correlated with the degree of demethylation. In addition, prolonged *in vitro* cultivation of either T^N^ or T^EMRA^ cells did not alter the methylation status of the *TBKBP1* DMR. Retroviral overexpression of TBKBP1 in CD8^+^ T cells resulted in TBK1 activation, as evidenced by an increased phosphorylation of TBK1. Furthermore, CMV-specific T cells overexpressing TBKBP1 showed enhanced cytotoxicity against CMV-infected target cells along with augmented pro-inflammatory cytokine production. In summary, our results suggest that targeted demethylation of the *TBKBP1* DMR or overexpression of TBKBP1 in CD8^+^ T cells might improve the current adoptive T cell therapy for the prevention of CMV infection and relapse.

## MATERIAL AND METHODS

### Human donors

All human samples were obtained from male healthy donors at the Institute of Transfusion Medicine and Transplant Engineering, Hannover Medical School (MHH). Written informed consent was obtained from all donors (MHH ethics committee votes 3639-2017, 9001_BO-K, 9255_BO_K_2020). For the phenotypic characterisation and isolation of T(CMV) cells, CMV-seropositive HLA-typed donors showing >0.5% CMV phosphoprotein 65 (pp65)-specific T cells upon antigen-specific restimulation (38,39) were selected. The dynamic virus reduction assay and immunoblotting experiments were performed using cells from male CMV-seronegative, human leukocyte antigen-A2 (HLA-A*02) negative donors. All remaining experiments were conducted using cells from male CMV-seropositive donors regardless of their HLA-type. CMV-serological testing and HLA-typing were conducted at the Institute of Transfusion Medicine and Transplant Engineering, MHH.

### Human sampling and cell isolation

For the phenotypic characterisation and whole genome bisulfite sequencing (WGBS) of T(CMV) cells, PBMCs were isolated from residual blood samples from disposable kits used during routine platelet apheresis at the Institute of Transfusion Medicine and Transplant Engineering, MHH, via discontinuous gradient centrifugation and immediately processed. For all remaining experiments, PBMCs were isolated from Leukocyte Reduction System (LRS) cones using Ficoll-based density gradient centrifugation (Lymphoprep; STEMCELL Technologies) and SepMate tubes (STEMCELL technologies). PBMCs were cryopreserved in FCS containing 10% DMSO. One day prior to the experiment, cryopreserved PBMCs were thawed, rinsed with excess TexMACS medium (Miltenyi Biotec) to remove DMSO, and rested overnight with TexMACS medium supplemented with 100 IU/ml of recombinant human IL-2 (Miltenyi Biotec).

### Antibodies for flow cytometry

All monoclonal antibodies used for immunophenotyping and cell sorting were purchased from BioLegend and BD Biosciences. Characterisation of T(CMV) cells was performed using anti-CD3 BV510, anti-CD8 FITC, anti-IFN-γ PE, anti-CD45RA PerCP/Cy5.5, and anti-CD62L AF647. For WGBS, T(CMV) cells and control T cell subsets were isolated using anti-CD3 AF488, anti-CD4 APC-Cy7, anti-CD8 APC, anti-CD27 BV510, anti-CD28 PE-Dazzle, anti-CD45RA PerCP/Cy5.5, anti-KLRG1 PE/Cy7, anti-CCR7 BV421, and anti-CX3CR1 BV650. For pyrosequencing, quantitative real-time-PCR (qRT-PCR), and *in vitro* cultivation experiments, CD8^+^ T cell subsets were isolated using anti-CD3 AF488, anti-CD8 APC, anti-CD28 PE Texas Red, anti-CD45RA PerCP/Cy5.5, anti-CD62L PE-Cy7, anti-CD95 PE, and anti-CCR7 BV421. For the isolation of retrovirally-transduced CMV-specific CD8^+^ T cells, anti-CD8 APC and anti-mouse TCR APC-Cy7 along with LIVE/DEAD® Fixable Blue Dead Cell dye were used (ThermoFischer Scientific).

### Flow cytometry

PBMCs were prepared for flow cytometry assay as described previously (40). Briefly, for surface staining, cells were resuspended in staining buffer (PBS and 0.5% BSA), and single-cell suspensions were labelled with antibodies for 30 minutes at 4°C, while chemokine receptor staining was carried out at 37°C for 30 minutes. Following surface staining, cells were washed and resuspended in staining buffer. Samples were acquired on a FACSCanto10c or LSR-II flow cytometer (BD Biosciences) and analysed using BD FACSDiva Software version 8.0.1 (BD Biosciences) and Flowjo v10 (BD Biosciences). Cell sorting of CD8^+^ T cell subsets and transduced cells was performed on a BD FACS ARIA II SORP (BD Biosciences).

### Phenotypic characterisation of T(CMV) cells

For the phenotypic characterisation of T(CMV) cells, CD8^+^ T cells were enriched from PBMCs using the untouched CD8^+^ T Cell Isolation Kit (Miltenyi Biotec) according to the manufacturer’s instructions. Next, the enriched CD8^+^ T cells were stimulated with the CMVpp65 overlapping peptide pool PepTivator® CMVpp65 (Miltenyi Biotec) at a concentration of 1 µg of each peptide/ml for 4 hours. Subsequently, IFN-γ-secreting cells were detected using IFN-γ Secretion Assay/Detection Kit (PE) (Miltenyi Biotec) according to the manufacturer’s instructions and subsequently phenotypically characterised by flow cytometry.

### Isolation of CD8^+^ T cell subsets for WGBS, pyrosequencing, qRT-PCR, and *in vitro* cultivation experiments

For the isolation of CD8^+^ T cell subsets for WGBS, PBMCs were first enriched for CD8^+^ T cells and subsequently stimulated with CMVpp65 overlapping peptide pool as described above. After the detection of IFN-γ-secreting cells, cells were stained and T(CMV) cells were isolated by flow cytometry-based cell sorting as CD3^+^CD4^−^CD8^+^IFN-γ^+^ cells, while bulk memory CD8^+^ T cells (T_mem_; CD3^+^CD4^−^CD8^+^CCR7^−^CD28^high^CD27^+^CD45RA^−^) and naive CD8^+^ T cells (T_N_; CD3^+^CD4^−^CD8^+^CCR7^+^CD28^int^KLRG1^−^CX3CR1^−^CD45RA^high^) were sorted as controls.

For pyrosequencing, qRT-PCR and *in vitro* cultivation experiments, human CD8^+^ T cell subsets were isolated from cryopreserved PBMCs. First, CD8^+^ T cells were enriched using human anti-CD8 MicroBeads and the automated magnetic activated cell sorting (autoMACS) system (both Miltenyi Biotec) according to the manufacturer’s instructions. Subsequently, the CD8^+^ T cells were stained and sorted into T_N_ (CD45RA^+^CCR7^+^), T_SCM_ (CD45RA^−^CCR7^+^CD28^+^CD62L^+^CD95^+^), T_CM_ (CD45RA^−^CCR7^+^), T_EM_ (CD45RA^−^CCR7^−^), and T_EMRA_ (CD45RA^+^CCR7^−^) CD8^+^CD3^+^ T cell subsets by flow cytometry.

### Whole-genome bisulfite sequencing (WGBS)

For WGBS, sorted T_N_, T_mem_, and T(CMV) cells from the blood of various donors were used to prepare genomic DNA with the DNeasy Blood & Tissue Kit (Qiagen). Approximately 50 ng genomic double-stranded DNA per donor was converted with sodium bisulfite using the EZ DNA Methylation-Lightning Kit (Zymo Research) and fragmented by sonication (Covaris S220, 10% duty cycle, 175W peak incident power, intensity 5, 200 cycles per burst, 120 seconds). The fragmented DNA served as input for the Accel-NGS Methyl-Seq DNA Library Kit (Swift Biosciences) and resulted in libraries that were sequenced on an Illumina NovaSeq 6000 with depths between 216 and 298 million paired end reads (2 x 100 bp).

Sequencing data were processed using the nf-core/methylseq pipeline (version 2.2.0), using default parameters, genome assembly GRCh38 and --accel=true (41) (software freely accessible at http://doi.org/10.5281/zenodo.1343417). Briefly, the pipeline employs FASTQC (version 0.11.9), trimgalore (version 0.6.7) and bismark (version 0.24.0) (42) for read-level quality control, adapter trimming, bisulfite-aware alignment, and cytosine-level DNA methylation quantification.

Methylation calls were further processed using RnBeads (version 2.17.0) (43) using a minimum per CpG coverage of 2 and removing CpGs that were covered in less than 50% of the samples, that overlapped annotated SNPs, or that were located on sex chromosomes. Genome-wide methylation values were computed by coverage weighted aggregation of CpG-level methylation values across samples in groups for 1 kb and 10 kb tiling windows. Dimensionality reduction using principal component analysis (PCA) was performed on those 50,000 1 kb tiling windows that exhibited the highest variance in aggregate methylation levels across the dataset. Pairwise DMRs between the groups were identified using Bsmooth/BSseq (version 1.30.0) (44). Briefly, we applied the Bsmooth() command with default parameters on unfiltered CpG methylation calls. Subsequently, only CpGs with a coverage of at least 5 in two or more samples per group were retained. Based on these filtered CpGs, we computed *t*-statistics and DMRs using BSmooth.tstat (…, estimate.var = "same", local.correct = TRUE) and dmrFinder (…, qcutoff=c(0.01,0.99), maxGap=200). Gene relations to nearest genes were annotated using GENCODE (version 44). Finally, DMRs containing at least 3 CpGs were selected using an absolute mean methylation level difference of 25%.

### *In vitro* cultivation of CD8^+^ T cell subsets

For *in vitro* cultivation, sorted T_N_ and T_EMRA_ cells were stimulated with plate-bound anti-human CD3 (OKT3; 1 μg/ml; BioLegend) and anti-human CD28 (CD28.2; 0.5 μg/ml; BioLegend) antibodies and cultured in TexMACS medium supplemented with 100 IU/ml human IL-2 at 37°C with 5% CO_2_. Every 5 days during the 30-day culture period, cells were harvested and washed, and an aliquot was taken for pyrosequencing and phenotypic characterisation by flow cytometry. The remaining cells were re-stimulated with plate-bound anti-human CD3 and anti-human CD28 (both 0.5 µg/ml) antibodies and cultured in fresh TexMACS medium supplemented with 100 IU/ml human IL-2.

### Pyrosequencing of the *TBKBP1* DMR

Genomic DNA from *ex vivo* isolated CD8^+^ T cell subsets or *in vitro* cultured cells was extracted using the DNeasy Blood & Tissue Kits (Qiagen) and subsequently bisulfite-converted by using the EZ DNA Methylation-Lightning Kit (Zymo Research) according to the manufacturers’ recommendations. *TBKBP1* DMR was pyrosequenced as described previously (45). For amplification of the *TBKBP1* DMR and subsequent pyrosequencing, we used the following primers:

‘forward’ 5’-TATTTAAGTTTGGGTGATAGAGTAAGAT-3’

‘reverse’ 5’-Bio-CCCAACCCTCAAAAATATAATATCT-3’

‘sequencing 1’ 5’-GGTGGTGTATGTTTGTAAT-3’

‘sequencing 2’ 5’-GGTTGAGGTGAGTTAAGAT-3’

### Quantitative RT-PCR of *TBKBP1*

Total RNA was purified from isolated CD8^+^ T cell subsets using the RNeasy Mini Kit (Qiagen), spectrometrically quantified (DeNovix), and transcribed into cDNA using Transcriptor First Strand cDNA Synthesis Kit (Roche). SYBR green master mix (Roche), cDNA, and TBKBP1 primers (‘forward’ 5’-TCGCTCTCATCACTGCCTAC-3’; ‘reverse’ 5’-GGGTACTTGATTTCATAGACTTTA-3’) were used in a qRT-PCR on a LightCycler 480 II (Roche). The data were analysed with LightCycler® 96 SW 1.1 (Roche). All procedures were performed according to the manufacturers’ recommendations.

### Retroviral transduction

For retroviral overexpression of TBKBP1, an expression cassette consisting of *TBKBP1* cDNA, porcine teschovirus-1 2A element (P2A), and mCherry cDNA was inserted into the pMP71 plasmid by gene synthesis (TWIST Bioscience). pMP71 containing only mCherry cDNA served as empty vector (EV) control. For the dynamic virus reduction assay, cells were not only retrovirally transduced with the TBKBP1 overexpression construct (or the EV control), but also with a pMP71-based plasmid coding for mTCR 5-2, a previously reported CMV-specific, HLA-A*02:01-restricted high-avidity T cell receptor (TCR) recognising the CMVpp65-derived peptide NLVPMVATV and containing a murine constant region (mTCR) for the isolation of successfully transduced cells (46).

Virus transduction of human PBMCs was performed as previously described (47). In brief, the virus-packaging RD114 cell line was seeded in a 6-well plate at a density of 1.5 x 10^6^ cells/well 1 day prior to transfection. The next day, pMP71 vectors were transfected using calcium phosphate precipitation. Thereto, complete DMEM medium (Gibco) supplemented with 10% FBS (Gibco), 1% sodium pyruvate (Biochrom), and 1% HEPES (Sigma-Aldrich) was replaced 1 hour before transfection with complete medium containing 25 μM chloroquine diphosphate salt (Sigma-Aldrich). The plasmid-containing buffer was prepared by adding 18 μg of vector DNA and 15 μl of 2.5 M CaCl_2_ (Applichem) to a final volume of 150 μl with water, which was then mixed with transfection buffer containing 280 mM sodium chloride, 42 mM HEPES, 3.5 mM disodium hydrogen phosphate, and 10 mM potassium chloride at 1:1 ratio by vortexing and subsequently incubated for 30 minutes at room temperature. The mixture was slowly added to RD114 cells (150 μl/well) in a dropwise manner and incubated for 15 hours at 37°C with 5% CO_2_, followed by medium replacement with 3 ml complete DMEM medium. The supernatant containing retroviral particles was harvested 3 days later. For transduction, PBMCs were stimulated for 72 hours with plate-bound anti-human CD3 and anti-human CD28 (both 1 μg/ml). Additionally, retronectin-treated plates were prepared by coating with 12.5 μg/ml retronectin (TaKaRa) in PBS (Gibco) at 4°C overnight in a non-treated 24-well plate. Unbound retronectin was aspirated and plates were blocked with 2% BSA (Sigma-Aldrich) in PBS for 20 minutes at 37°C followed by washing twice with PBS. Supernatant containing retroviral particles was added to retronectin-coated plates followed by centrifugation at 2,000xg for 2 hours at 32°C. Following centrifugation, virus supernatant was aspirated and preactivated PBMCs (0.5-0.8 x 10^6^/well) in TexMACS medium supplemented with 100 IU/ml IL-2 and 6 ng/ml polybrene (Sigma-Aldrich) were added to each well. After spinoculating the cells onto virus-coated plates (1,000xg for 10 minutes at 32°C), the cells were incubated at 37°C with 5% CO_2_ for 24 hours. After 24 hours, transduced human PBMCs were washed and resuspended in TexMACS medium supplemented with IL-2. Flow cytometry analysis was performed 5 days after spinoculation to determine transfection efficiency. Successfully transduced CD8^+^mCherry^+^ T cells (TBKBP1 overexpression or EV control) or CD8^+^mTCR^+^mCherry^+^ T cells (simultaneous overexpression of mTCR 5-2 and TBKBP1 or EV control) were sorted using flow cytometry and used for subsequent experiments.

### Cell stimulation and immunoblotting

For immunoblotting experiments, CD8^+^ T cells overexpressing TBKBP1 (or EV controls) were generated and sorted as described above. Next, cells were serum-starved overnight and subsequently incubated with soluble 2 μg/ml anti-CD3 and anti-CD28 mAbs for 40 minutes on ice. Antibody-coated cells were then cross-linked with 4 μg/ml AffiniPure goat-anti-mouse IgG+IgM (H+L; Jackson ImmunoResearch) and incubated for 15 minutes at 37°C. Unstimulated controls were kept on ice. To stop the stimulation, excess ice-cold PBS was added and the cells were centrifuged at a pre-cooled temperature. Pellets were snap-frozen and stored at -80°C. For preparing cell lysates for SDS-PAGE and immunoblotting, cell pellets were resuspended in 40 µl of 2x SDS loading buffer (0.125 M Tris-HCl, pH6.8, 4% SDS, 20% glycerol, 0.02% bromphenol blue, 10% β-mercaptoethanol), heated at 95°C for 5 minutes, and cell debris was pelleted with 14,000xg at room temperature (RT) for 5 minutes. Per lane, 20 µl lysate was analysed on 10% polyacrylamide gels containing 0.1% SDS in Tris-glycine running buffer with 0.1% SDS at RT. Proteins were blotted on polyvinylidene difluoride membranes (Amersham Hybond P 0.2 µm) in a wet blot system in Tris-glycine transfer buffer containing 0.05% SDS and 20% methanol for 1 hour at 350 mA at 4°C. Membranes were subsequently blocked in 5% BSA in TBS with 0.1% Tween 20 for 1 hour at RT and incubated with anti-phospho-TBK1/NAK (Ser 172), anti-SINTBAD (TBKBP1; D1A5), or anti-GAPDH (14C10) primary antibodies (all from Cell Signaling Technology) at 4°C overnight. Incubation with HRP-coupled anti-rabbit secondary antibody (Dianova) was carried out for 1 hour at RT. Membranes were developed with the Pierce ECL or SuperSignal substrate (Thermo Fisher Scientific) and a ChemoStar ECL Imager (INTAS). Quantitative analysis was performed with ImageJ software (version 1.52g, National Institute of Health). Membranes were stripped with Restore PLUS Western Blot Stripping Buffer (ThermoFisher Scientific) for 15 minutes at room temperature, blocked, incubated with an anti-TBK1/NAK (D1B4) primary antibody (Cell Signaling Technology) and anti-rabbit secondary antibody, imaged, and analysed as described above.

### Dynamic virus reduction assay

For the dynamic virus reduction assay, CD8^+^ T cells simultaneously overexpressing mTCR 5-2 and TBKBP1 (or EV controls) were generated and sorted as described above. One day prior to infection, MRC-5 cells were seeded in 96-well plates at a density of 20,000 cells/well and cultured in Eagle’s minimum essential medium with 10% FCS, 1 mM sodium pyruvate, and 1 mM glutamine. The following day, the MRC-5 cells were infected with a reporter CMV (TB40/BAC4 HCMV^3F^) that was generated on the background of the clinical isolate TB40-E (48). Briefly, after aspiration of the medium, the reporter CMV was added to the MRC-5 cells at a MOI of 0.1 and centrifuged at 700xg for 10 minutes at room temperature to enhance the infection and reduce infection heterogeneity. After centrifugation, the medium was immediately aspirated and the cells were washed with PBS. After washing, sorted mTCR^+^mCherry^+^ CD8^+^ T cells were added to the infected MRC-5 cells in a final volume of 200 μl medium at effector:target (E:T) ratios of 0.5:1 and 1:1. Coculture medium was composed of CTS™ OpTmizer™ T-cell expansion basal medium (ThermoFischer Scientific) supplemented with CTS™ Immune Cell SR (ThermoFischer Scientific) and 300 IU/ml IL-2. To analyse the virus reduction capacity of the T cells, the mNeonGreen signals from infected MRC-5 cells were monitored for 1 week using the Incucyte™ Live-Cell analysis system (Sartorius). Additionally, the confluence of cells was determined by the phase detector and images were analysed by the Incucyte™ Analysis Software. Data were exported to Prism 9.4.0 for graphical representation.

### Cytokine multiplex assay

For measuring cytokines, 50 μl culture supernatants were harvested after 36 hours of the co-culture from the dynamic virus reduction assay. Supernatants were centrifuged at 500xg for 15 minutes at 4°C and kept at -80°C. For cytokine measurement, 50 μl thawed supernatant was diluted 1:1 with supplied assay buffer. Cytokines were measured using the MILLIPLEX® Human CD8^+^ T Cell Magnetic Bead Panel Premixed 17 Plex-Immunology Multiplex Assay (Millipore) according to the manufacturer’s recommendations and analysis was done using the Bio-Plex Manager 6.1 software based on the standard curve.

### Statistical analysis

Prism 9.4.0 software was utilized for statistical analysis. The normal distribution of data points was checked using the Shapiro-Wilk normality test. A paired two-tailed parametric student’s *t* test was conducted on samples that passed the normality test, and a non-parametric Wilcoxon matched-pairs signed rank test was performed on samples that failed in normality test. All data are presented as mean or mean ± SD, and *p*-values ≤ 0.05 deemed significant (* p ≤ 0.05; ** p ≤ 0.01; *** p ≤ 0.001; **** p ≤ 0.0001; ns; not significant).

### Data availability

Sequencing data and methylation levels reported in this paper were uploaded to GEO under accession number GSE245832. Identified DMRs are listed in Supplementary Table 1.

## RESULTS

### T(CMV) cells exhibit a unique epigenetic signature

The differentiation of naive murine CD8^+^ T cells into the different types of memory and effector T cells is associated with specific changes at the transcriptional and epigenetic level (23,24,29,49,50). Studies on human CD8^+^ T cell memory formation similarly identified changes in DNA methylation patterns in the cytotoxicity-related genes *IFNG*, *GZMB,* and *PRF1* (51,52). However, the global changes in DNA methylation patterns during the pathogen-driven formation of antigen-specific memory CD8^+^ T cells are only incompletely understood. Therefore, we isolated T(CMV) cells from 5 different CMV-seropositive donors to unravel their specific DNA methylation patterns. T(CMV) cells were identified as IFN-γ-secreting cells after restimulation with an overlapping peptide pool of CMVpp65 (65 kDa lower matrix phosphoprotein), which is the main component of the enveloped subviral particle and an immunodominant target of T cell responses to CMV (53) (Supplementary Figure 1). An initial flow cytometric characterisation revealed that the vast majority of these IFN-γ^+^ T(CMV) cells showed a CD45RA^+^CD62L^−^ T_EMRA_ cell phenotype (Supplementary Figure 1), in line with recently published data (54,55). For whole-genome bisulfite sequencing (WGBS), T(CMV) cells were isolated by flow cytometry-based cell sorting as CD3^+^CD4^−^CD8^+^IFN-γ^+^ cells, while bulk memory CD8^+^ T cells (T_mem_; CD3^+^CD4^−^CD8^+^CCR7^−^CD28^high^CD27^+^CD45RA^−^; n=5) and naive CD8^+^ T cells (T_N_; CD3^+^CD4^−^CD8^+^CCR7^+^CD28^int^KLRG1^−^CX3CR1^−^CD45RA^high^; n=4) from the same donors served as controls (Supplementary Figure 2). For each sorted sample, genomic DNA was isolated and the genome-wide DNA methylation profile was determined by WGBS. Global analysis of the DNA methylation data revealed a strong progressive loss of DNA methylation from T_N_ to both T_mem_ and T(CMV) cells (Figure 1A), in line with previously published data on human CD4^+^ memory T cells (56). Next, we identified DMRs in pairwise comparisons of all T cell subsets. We found the highest numbers of DMRs between T_mem_ vs. T_N_ cells (11,357) and T(CMV) vs. T_N_ cells (13,011), while a rather low number of only 104 DMRs were identified in the comparison of T(CMV) vs. T_mem_ cells (Supplementary Table 1). Accordingly, a PCA based on genome-wide DNA methylation levels revealed a close sample group relationship between T(CMV) and T_mem_ cells, whereas T_N_ cells were placed separately along the main principal component 1 (PC1) (Figure 1B). This finding was confirmed by hierarchical clustering of individual samples, which further demonstrated that the majority of DMRs showed a consistent methylation status in T(CMV) and T_mem_ cells, while being distinct to the largely methylated state of most DMRs in T_N_ cells (Figure 1C). Yet, it is important to note that all T(CMV) and T_mem_ cell samples clustered group-wise, indicating distinct methylation signatures between these cell types. The majority of DMRs across all pair-wise population comparisons were located in gene bodies and intergenic regions, while a smaller fraction was mapped to promoters (Figure 1D). In line with previous findings (33,57), we observed that many genes involved in the CD8^+^ T cell effector program contain demethylated regions in both T(CMV) and T_mem_ cells, but are methylated in T_N_ cells, including genes coding for molecules regulating T cell-mediated cytotoxicity (*GZMA*, *GZMB*, *GZMK*, *IFNG*), actin cytoskeletal organization (*ACTA2*, *ACTB*, *ACTN4*, *DOCK2*), integrin-dependent cell adhesion (*ITGB1*, *ITGB1BP1*, *ITGA5*), chemokine-mediated signalling (*CCL1*, *CCL4*, *CCL5*, *CCR4*, *CCR5*), TGFβ-mediated signalling (*SMAD2*, *SMAD3*, *SMAD7*), and WNT/β-catenin signalling (*APC2*, *CTR9*, *CYLD*, *WNT11*, *WNT7B*) (Supplementary Table 1). Taken together, the WGBS study demonstrated substantial changes of DNA methylation patterns during the differentiation of T_N_ into T(CMV) cells.

**Figure 1:**
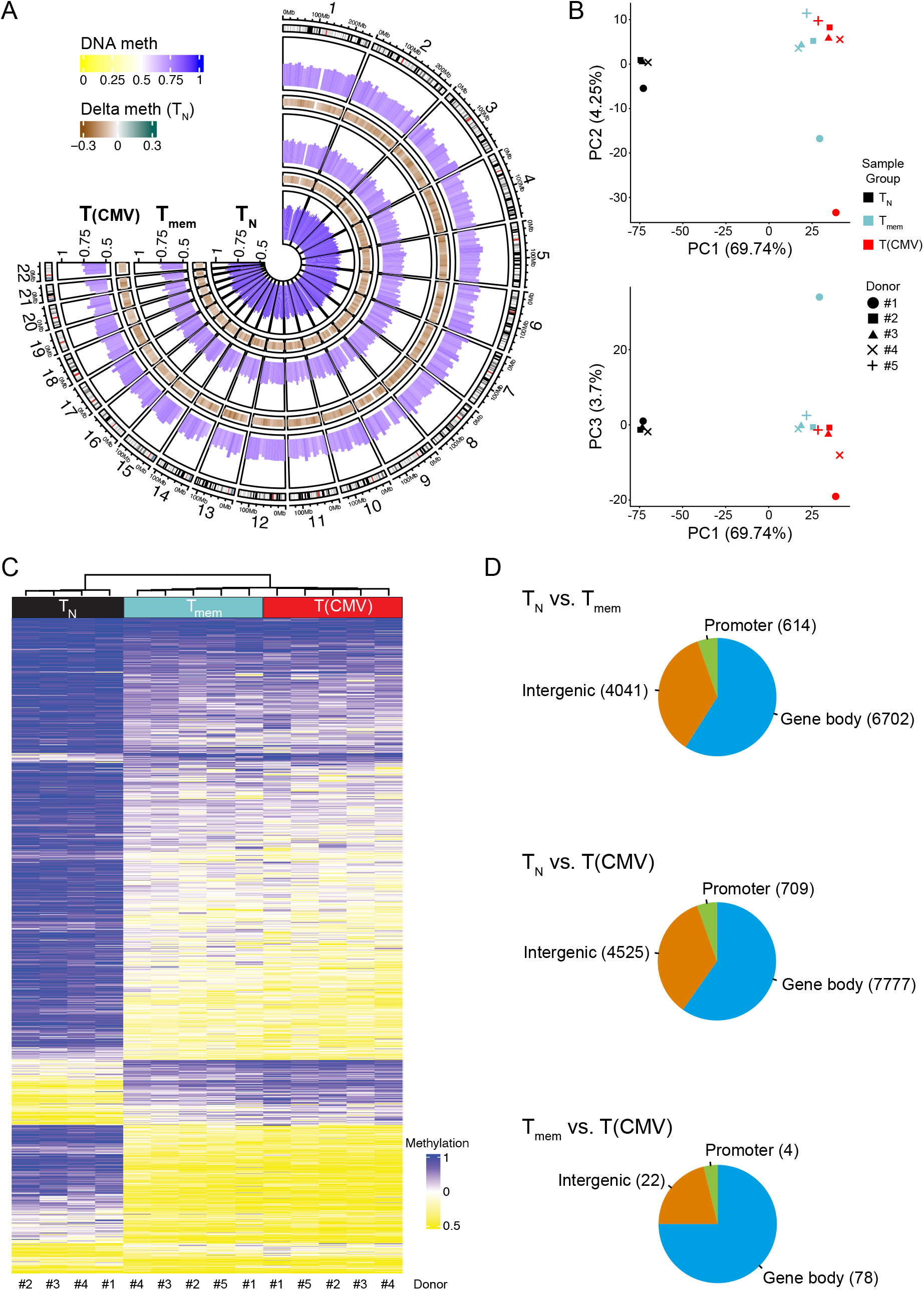
Genome-wide methylation profiling of T(CMV) cells. CD8^+^ T cell subsets including T_N_ (n=4), T_mem_ (n=5), and T(CMV) cells (n=5) were sorted using flow cytometry. From each sample, genomic DNA was isolated and converted by bisulfite treatment, followed by WGBS. (A) Circos plot showing DNA methylation levels for T_N_, T_mem_, and T(CMV) cells across the whole genome (in 10 kb tiling windows, aggregated all donors). CpG methylation levels are represented as histogram tracks across the genome based on the sample. High levels of methylation are indicated by dark blue, while low levels of methylation are indicated by light yellow. Brown-teal colour-coding between the bar tracks indicates differences in methylation levels relative to T_N_ cells. (B) Principal Component Analysis (PCA) per cell type and donor samples based on 50,000 highly variable 1 kb tiling regions. The percent of explained variance for each component is denoted in the axis labels. (C) Hierarchical clustering of 15,598 non-overlapping DMRs identified in pair-wise comparisons of methylomes from T_N_, T_mem_, and T(CMV) cells. The dendrogram on top corresponds to hierarchical clustering of the samples. The colour-code illustrates the mean methylation levels of the DMRs as indicated (yellow: methylation level = 0%, blue: methylation level = 100%) (D) Pie charts indicating the position of the pairwise DMRs identified in indicated group-wise comparisons relative to annotated genes. Numbers in parentheses show the number of DMRs in intergenic, promoter, or gene body regions according to their genomic position.

Next, we focused our analysis on the 104 DMRs that were identified in the comparison of T(CMV) vs. T_mem_ cells. A small fraction (25 DMRs) was also detected in the comparison of T_mem_ vs. T_N_ cells, and showed a more pronounced demethylation in T(CMV) cells (Supplementary Table 1). However, the majority (79 DMRs) were only detected in the comparison of T(CMV) vs. T_mem_ cells, and thus determine the unique epigenetic signature of T(CMV) cells. It is important to note that in a few cases more than 1 DMR was found in proximity to a given gene, resulting in 71 unique genes showing differential DNA methylation patterns between T(CMV) and T_mem_ cells (Supplementary Table 1). Some of the top demethylated DMRs in T(CMV) cells, identified in the pairwise comparison with T_mem_ cells, were associated to genes with known roles in differentiation and cytotoxic function of CD8^+^ T cells, including *CLDND2*, *KLRD1*, *TBX21*, *ZEB2*, and *S1PR5* (Figure 2). In addition, a number of DMRs being selectively demethylated in T(CMV) cells were associated to genes for which a role in CD8^+^ T cells has not been described yet, namely *FGR*, *LINC01871*, *TMEM14C*, *MAD1L1*, *PIBF1*, *LMF1*, and *TBKBP1*. TBK-binding protein 1 (TBKBP1) is an adaptor protein that can interact with TANK-binding kinase 1 (TBK1) and promote its activation (58,59). TBK1 is well known to mediate the activation of interferon regulatory factors (IRFs), namely IRF3 and IRF7, as well as NF-κB signalling to regulate pro-inflammatory cytokine production and the activation of innate immunity (60–66). The transcription signature of CD8^+^ T_EMRA_ cells can be regulated by several innate immune-related genes, including natural killer (NK) cell activation mediators and multiple NK receptors (33). These findings prompted us to study the role of TBKBP1 in CD8^+^ T cells in more detail.

**Figure 2:**
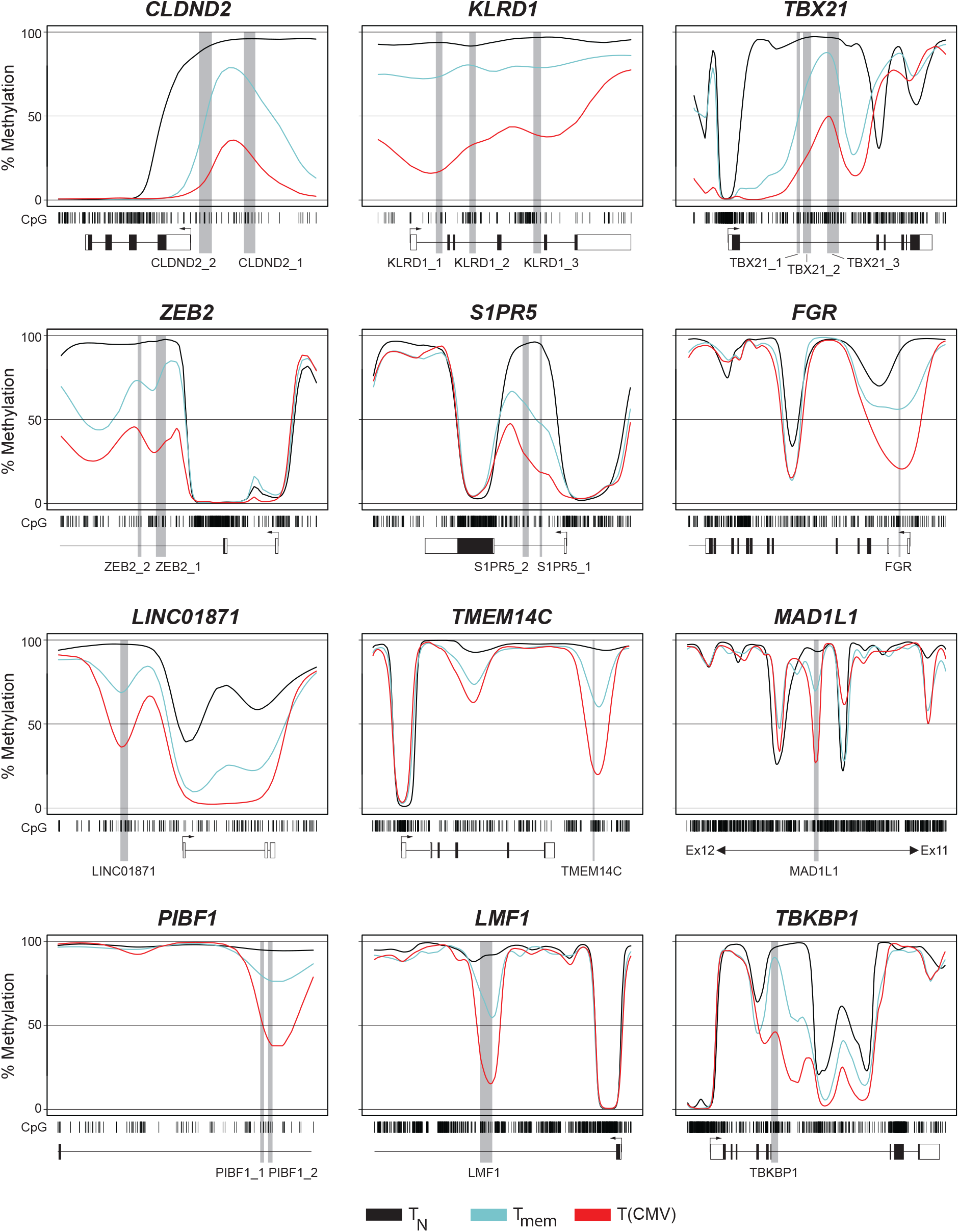
Methylation profiles of gene loci associated with top DMRs from epigenetic signature of T(CMV) cells. Out of the 71 genes showing unique differential DNA methylation patterns between T(CMV) and T_mem_ cells the top 12 DMR-associated gene loci were selected. For each gene, CpG motifs (barcodes), DMRs (light grey boxes), and exons of the surrounding gene body (dark grey boxes, transcriptional start site indicated by arrows) are displayed. Coloured lines illustrate methylation values ranging from 0-100% of T_N_ (black), T_mem_ (cyan), and T(CMV) cells (red) in a linear manner. Mean values from all donors per CD8^+^ T cell subset are depicted.

### Demethylation of the *TBKBP1* DMR is associated with higher *TBKBP1* gene expression in effector memory subsets

To characterise the methylation status of the newly identified *TBKBP1* DMR within major human CD8^+^ T cell subpopulations, we isolated genomic DNA from sorted CD45RA^+^CCR7^+^ T_N_, CD45RA^+^CCR7^−^ T_EMRA_, CD45RA^−^CCR7^−^ T_EM_, CD45RA^−^CCR7^+^ T_CM_, and CD45RA^−^CCR7^+^CD28^+^CD62L^+^CD95^+^ T_SCM_ cells isolated from peripheral blood and subjected them to pyrosequencing analysis. Analysis of the *TBKBP1* DMR showed a pronounced demethylation pattern in T_EMRA_ cells and a partial demethylation in T_EM_ cells, while T_CM_, T_SCM_ and particularly T_N_ cells were largely methylated at these sites (Figure 3A). Since it has been previously reported that CpG methylation levels around the promoter or first intron/exon region of a specific gene are inversely correlated with gene expression (67–69), we next asked if the strong demethylation of the *TBKBP1* DMR in T_EMRA_ cells is accompanied by a high *TBKBP1* expression. Indeed, analysis of *TBKBP1* gene expression revealed a high expression in T_EMRA_ cells, an intermediate expression in T_EM_ cells, a low expression in T_CM_ and T_SCM_ cells, while *TBKBP1* expression was below the detection level in T_N_ cells (Figure 3B). Hence, we could observe a clear inverse correlation between DNA methylation rates and *TBKBP1* gene expression levels (r = -0.7575; *p*-value = <0.00010) (Figure 3C), indicating a DMR-mediated transcriptional control of the *TBKBP1* gene locus.

**Figure 3.**
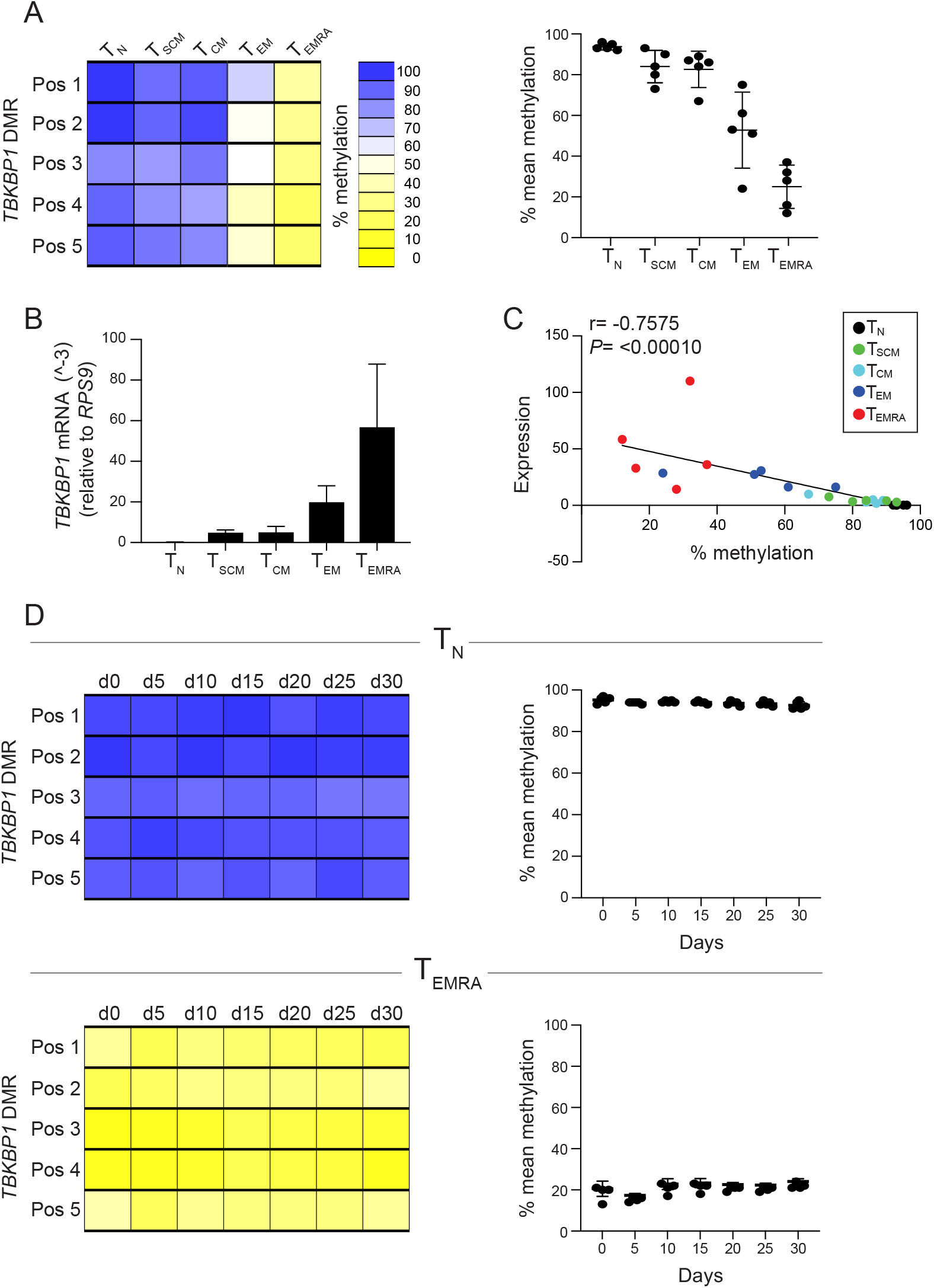
Demethylation of *TBKBP1* DMR correlates with increased *TBKBP1* expression in T_EMRA_ and T_EM_ CD8^+^ T cell subsets and is stably maintained upon *in vitro* culture. Indicated CD8^+^ T cell subsets were isolated from CMV-seropositive healthy donors and genomic DNA as well as RNA were isolated from sorted samples. Bisulfite-converted genomic DNA was subjected to pyrosequencing using primers targeting the *TBKBP1* DMR and RNA was transcribed into cDNA to determine *TBKBP1* expression levels by qRT-PCR. (A) Methylation profiles of the *TBKBP1* DMR in indicated CD8^+^ T cell subsets. (Left) The methylation values from 1 representative donor were translated into a colour-code according to the scale ranging from yellow (0% methylation) via white (50% methylation) to blue (100% methylation), the position of the CpG motifs is depicted, and each rectangle represents the methylation value of a single CpG motif. (Right) The scatter plot shows the mean methylation level of all 5 CpG motifs of the *TBKBP1* DMR in indicated CD8^+^ T cell subsets from 5 independent donors. Each dot represents data from one donor and mean values±SD are depicted. (B) The bar plot shows the relative *TBKBP1* expression normalized to the housekeeping gene *RPS9* in indicated CD8^+^ T cell subsets. Mean values±SD are depicted (n=5). (C) Scatterplot and linear regression analysis show correlation of mean methylation of *TBKBP1* DMR with mean *TBKBP1* expression in indicated CD8^+^ T cell subsets from 5 donors (r, correlation coefficient; *p*, *p*-value). (D) CD8^+^ T_EMRA_ and T_N_ cells were isolated from CMV-seropositive healthy donors and cultured *in vitro* with plate-bound anti-CD3/CD28 antibodies in the presence of exogenous human IL-2 for up to 30 days. Every 5 days, samples from cultured T_N_ (top) and T_EMRA_ cells (bottom) were taken to analyse the *TBKBP1* DMR methylation status as described in panel A. The heatmaps are from a representative donor (left) and scatter plots summarize the mean methylation levels of all 5 CpG motifs of the *TBKBP1* DMR from 5 independent donors. Each dot represents data from one donor and mean values±SD are depicted.

During homeostatic proliferation, naive CD8^+^ T cells have been reported to undergo phenotypic alterations (70,71). This observation raised the question whether the methylated and demethylated states of the *TBKBP1* DMR in T_N_ and T_EMRA_ cells, respectively, remain stable during proliferation. To answer this question, we isolated CD8^+^ T_N_ and T_EMRA_ cells from peripheral blood of CMV-seropositive healthy donors and cultured them for up to 30 days in the presence of anti-CD3 and anti-CD28 antibodies plus IL-2. Every 5 days, an aliquot was taken for phenotypic characterisation by flow cytometry and pyrosequencing. Upon culture of T_N_ cells, we observed a transient downregulation of the homing receptor CCR7 reaching normal levels by day 20, and an acquisition of stem-like properties with the upregulation of CD62L and CD28 on day 15 (Supplementary Figure 3), as recently reported (52). In contrast, T_EMRA_ cells more stably maintained their CD45RA^+^CCR7^−^ phenotype throughout the culture. Pyrosequencing analyses revealed that the *TBKBP1* DMR remained fully methylated in cultured T_N_ cells and fully demethylated in cultured T_EMRA_ cells during the entire cultivation period (Figure 3D). Together, these findings suggest that the epigenetic status of the *TBKBP1* gene is rather stable and preserved in both cultured T_N_ and T_EMRA_ cells even after multiple rounds of TCR-triggered cell division.

### TBKBP1 overexpression promotes the phosphorylation of TBK1 in stimulated CD8^+^ T cells

Although it is well known that the kinase TBK1 can be activated by various signals, including TCR signalling (60,72), the role of its adaptor protein TBKBP1 in CD8^+^ T cells remains unidentified. Recently, TBKBP1 was reported to support the recruitment of TBK1 to protein kinase C-theta (PKCθ) in lung epithelial cells, thereby promoting TBK1 phosphorylation and activation (59) (Figure 4A). To investigate if the same mechanism applies in T cells, we retrovirally overexpressed TBKBP1 in CD8^+^ T cells and subsequently monitored TBK1 phosphorylation after short-term restimulation with anti-CD3 and anti-CD28 antibodies (Supplementary Figure 4A). Successfully transduced CD8^+^ T cells were sorted by flow cytometry (Supplementary Figure 4B) and expectedly, TBKBP1-overexpressing cells showed high *TBKBP1* mRNA expression levels when compared to EV-transduced control cells (Supplementary Figure 4C). Immunoblotting confirmed TBKBP1 protein expression in CD8^+^ T cells retrovirally transduced with the TBKBP1 overexpression construct, and no differences were observed between unstimulated and restimulated CD8^+^ T cells (Figure 4B). Similarly, neither TBKBP1 overexpression nor the short-term restimulation influenced TBK1 expression in CD8^+^ T cells (Figure 4C). However, the phosphorylation of TBK1 was significantly enhanced in TBKBP1-overexpressing CD8^+^ T cells, but only after short-term restimulation, while EV-transduced control cells did not show any increase in TBK1 phosphorylation when short-term restimulated were compared to unstimulated cells (Figure 4D, E). In summary, these results demonstrate that TBKBP1 expression promotes TBK1 phosphorylation in activated CD8^+^ T cells, likely resulting in the activation of further downstream signalling pathways.

**Figure 4:**
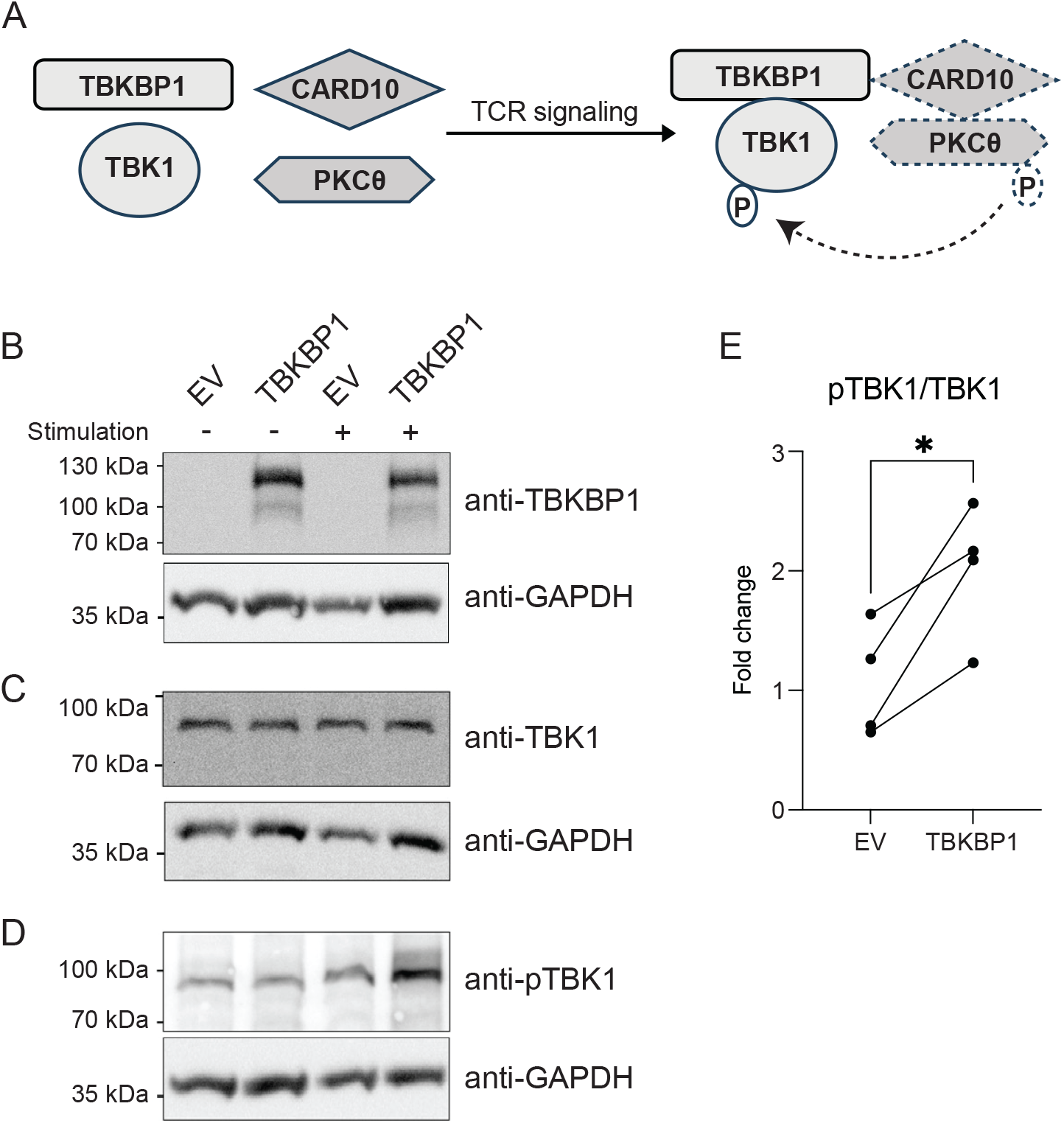
TBKBP1 overexpression in restimulated CD8^+^ T cells results in increased TBK1 phosphorylation. (A) The model illustrates how TCR signals might activate the phosphorylation of TBK1. Briefly, upon stimulation of T cells, TBKBP1 recruits TBK1 to PKCθ via CARD10, a scaffold protein. This association facilitates PKCθ to phosphorylate TBK1, an essential step in the activation of TBK1 and downstream signalling. This illustration is adapted from Zhu et al. (59). PBMCs obtained from CMV-seronegative healthy donors were stimulated with plate-bound anti-human CD3 and anti-human CD28 for 72 hours and subsequently retrovirally transduced with TBKBP1-expressing vectors or empty vector (EV) controls. Successfully transduced CD8^+^mCherry^+^ T cells were sorted by flow cytometry, serum-starved overnight, and subsequently stimulated with soluble anti-human CD3, anti-human CD28, and cross-linking antibodies, while unstimulated cells served as additional controls. From TBKBP1-overexpressing or EV-transduced CD8^+^ T cells both unstimulated (-) and stimulated (+) samples were subjected to immunoblotting to determine the expression of (B) TBKBP1, (C) TBK1, and (D) phosphorylated TBK1 (pTBK1). The analysis of GAPDH expression served as loading control. (E) Band intensities for pTBK1 and TBK1 were quantified and the fold change of TBK1 phosphorylation upon stimulation of EV-transduced or TBKBP1-overexpressing CD8^+^ T cells was determined by dividing the ratio of pTBK1/TBK1 band intensities of stimulated to the ratio of pTBK1/TBK1 band intensities of unstimulated samples. Data are from 4 independent donors analysed in 2 independent experiments. For statistical analyses, a paired two-tailed student’s *t* test was conducted with *, p ≤ 0.05.

### Ectopic expression of TBKBP1 augments the virus-reducing capacity of CD8^+^ T cells

After having demonstrated that TBKBP1 has an effect on TBK1 phosphorylation, we next sought to investigate its functional role in CD8^+^ T cells during an immune reaction. To this end, we utilized the dynamic virus reduction assay, in which fibroblasts (MRC-5 cells) infected with recombinant CMV expressing the reporter protein mNeonGreen (48) were co-cultured with CMV-specific CD8^+^ T cells, which were generated by retroviral overexpression of the HLA-A*02:01-restricted high-avidity TCR recognising the CMVpp65-derived peptide NLVPMVATV (mTCR 5-2) (46). To assess the functional role of TBKBP1, cells were additionally retrovirally transduced with the TBKBP1 overexpression construct, while EV-transduced cells served as control. Successfully transduced mTCR 5-2^+^mCherry^+^ CD8^+^ T cells were sorted by flow cytometry (Supplementary Figure 5) and added to the CMV-infected MRC-5 cells at different effector:target (E:T) ratios. The dynamic virus reduction was determined by measuring the expression of mNeonGreen using live cell imaging and representative microscopic images demonstrate the strong anti-viral and cytopathic effect of mTCR 5-2^+^ CD8^+^ T cells, irrespective of TBKBP1 overexpression, with an overall reduced number of viable mNeonGreen-expressing MRC-5 cells when compared to controls lacking T cells (Figure 5A). Hourly measurements of the reporter signal allowed a quantification of the dynamic virus reduction, and in cultures without T cells a continuous increase of the reporter signal was observed reaching a plateau around 12 hours with an only mild increase afterwards (Figure 5B, green curve). Upon addition of EV-transduced mTCR 5-2^+^ control CD8^+^ T cells, a marked decrease of the reporter signal was observed, starting at 12 hours and continuing until the end of the assay at 72 hours (Figure 5B, blue curve). This signal reduction was even more pronounced when TBKBP1-overexpressing mTCR 5-2^+^ CD8^+^ T cells were added, showing a significantly stronger neutralization of MRC-5-infected cells irrespective of the E:T ratio (Figure 5B, red curve). The improved virus reduction by TBKBP1-overexpressing CD8^+^ T cells was associated with significantly elevated levels of IFN-γ and Granzyme A, and a trend towards increased levels of Granzyme B, Perforin, MIP-1α (CCL3), and MIP-1β (CCL4) in supernatants of co-cultures with TBKBP1-overexpressing when compared to EV-transduced control CD8^+^ T cells (Figure 5C, Supplementary Figure 6). Collectively, these data demonstrate that ectopic expression of TBKBP1 in human CD8^+^ T cells results in the upregulation of pro-inflammatory cytokines and chemokines, facilitating a significantly enhanced virus neutralization capacity.

**Figure 5:**
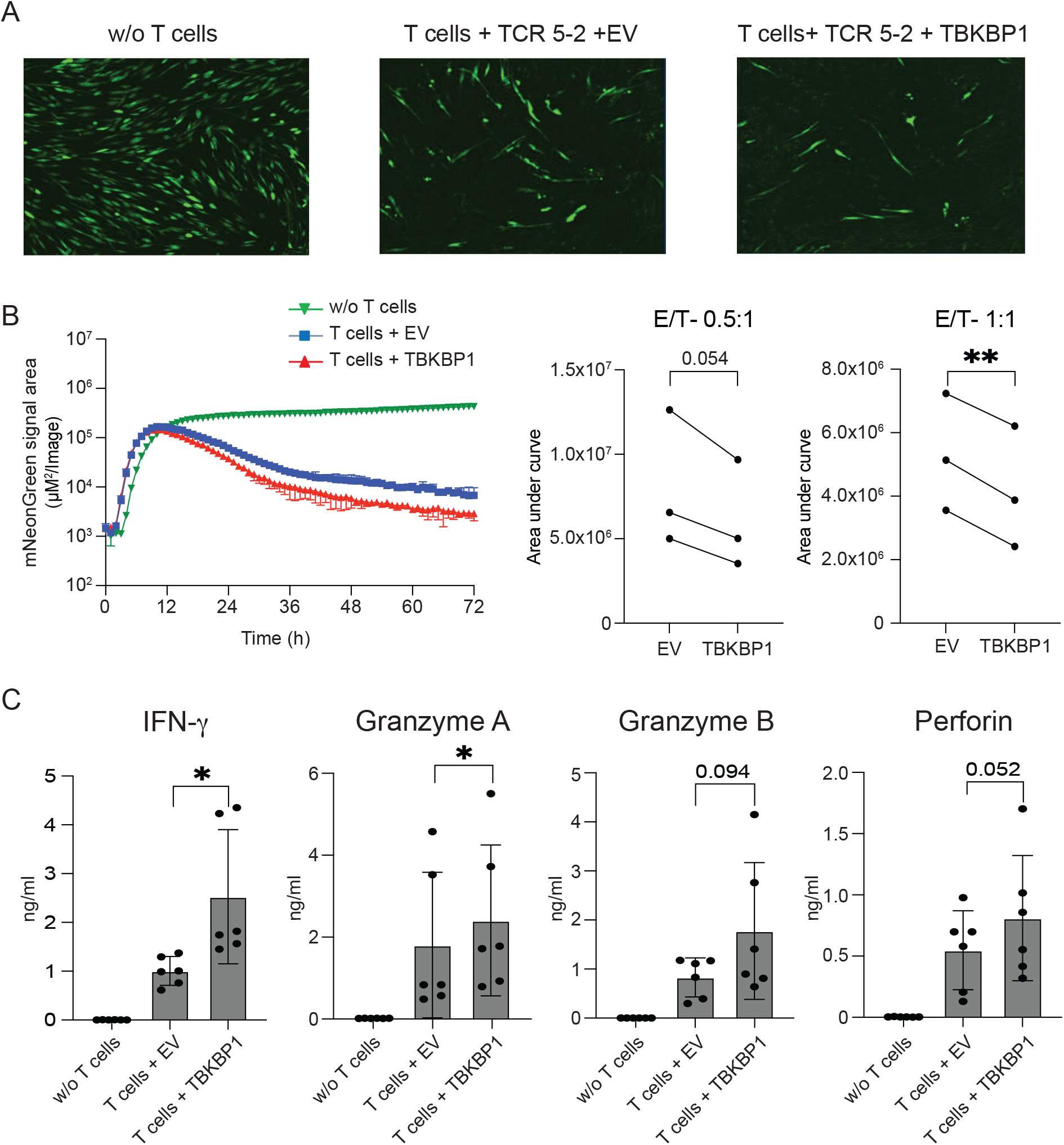
TBKBP1 overexpression enhances virus-reducing capacity of CD8^+^ T cells. PBMCs obtained from CMV-seronegative healthy donors were stimulated with plate-bound anti-human CD3 and anti-human CD28 for 72 hours and subsequently retrovirally transduced with both the high-avidity TCR 5-2 containing murine constant regions (mTCR) and directed against the CMV HLA-A*02-peptide NLVPMVATV and also TBKBP1-overexpressing vectors or empty vector (EV) controls. Successfully transduced mTCR 5-2^+^mCherry^+^ CD8^+^ T cells were sorted by flow cytometry and added at indicated effector:target (E:T) ratios to MRC 5 cells infected with recombinant CMV expressing the reporter protein mNeonGreen. Dynamic virus reduction was determined by measuring the expression of mNeonGreen using live cell imaging. (A) Representative microscopic images 48 hours after infection show the expression of mNeonGreen in CMV-infected MRC-5 cells in the absence of added T cells (w/o T cells), in the presence of EV-transduced mTCR 5-2^+^ CD8^+^ T cells (T cells + TCR 5-2 +EV) or in the presence of TBKBP1-overexpressing mTCR 5-2^+^ CD8^+^ T cells (T cells + TCR 5-2 +TBKBP1). T cells were added at a E:T ratio of 1:1. (B) The dynamic virus reduction was quantified by hourly measurements of the reporter signal in cultures in the absence of added T cells (w/o T cells, green curve), in the presence of EV-transduced mTCR 5-2^+^ CD8^+^ T cells (T cells + TCR 5-2 +EV, blue curve) or in the presence of TBKBP1-overexpressing mTCR 5-2^+^ CD8^+^ T cells (T cells + TCR 5-2 +TBKBP1, red curve). (Left) The graph depicts mean values±SD from 2 technical replicates of 1 representative out of 3 independent experiments. (Right) The line plots summarize data from 3 independent experiments by showing the average area under the curve (AUC) ±95 % CI for EV-transduced or TBKBP1-overexpressing mTCR 5-2^+^ CD8^+^ T cells cultured with CMV-infected MRC-5 at indicated E:T ratios. (C) 36 hours after infection, culture supernatants were harvested and cytokine profiles were determined from cultures of CMV-infected MRC-5 cells in the absence of added T cells (w/o T cells), in the presence of EV-transduced mTCR 5-2^+^ CD8^+^ T cells (T cells + EV, E:T=1:1) or in the presence of TBKBP1-overexpressing mTCR 5-2^+^ CD8^+^ T cells (T cells + TBKBP1, E:T=1:1). Data from 3 independent experiments with 2 technical replicates each are shown. For statistical analyses, a paired two-tailed student’s *t* test (leaving out “w/o T cell” group) was conducted with *, p ≤ 0.05 and **, p ≤ 0.01.

## DISCUSSION

Epigenetic processes, including DNA methylation, are well-known contributors to T cell fate specification upon CD8^+^ T cell differentiation (23–25). In the present study, we hypothesized that global DNA methylation analyses could provide insights into the epigenetic reprogramming of T(CMV) cells and also support the identification of key players involved in their effector functions. Utilizing WGBS, we observed substantial epigenetic alterations upon differentiation of CD8^+^ T_N_ into T_mem_ and T(CMV) cells, and the vast majority of identified DMRs were demethylated in both T_mem_ and T(CMV) when compared to T_N_ cells. With the final aim to identify an epigenetic signature of T(CMV) cells, we focused our analysis on the comparison of DNA methylation profiles of T_mem_ and T(CMV)_CMV_ cells, and could identify 79 regions being differentially methylated between T(CMV) and T_mem_ cells and not between T_mem_ and T_N_ cells, thus representing a unique epigenetic signature of T(CMV) cells. One of the epigenetic signature genes was *TBKBP1*, not described in the context of T cell-mediated immune responses so far. This encouraged us to investigate the specific role of TBKBP1 for the cytotoxic function of CD8^+^ T cells. We could validate the preferential demethylation of the *TBKBP1* DMR in T_EMRA_ cells, the dominant phenotype of T(CMV) cells, which also showed the highest *TBKBP1* mRNA expression levels. Furthermore, we observed a significant increase in TBK1 phosphorylation upon TCR stimulation in TBKBP1-overexpressing CD8^+^ T cells, suggesting that TBKBP1 is crucial for the optimal TCR-induced activation of TBK1 and downstream signalling pathways. Accordingly, TBKBP1 overexpression resulted in an increased production of pro-inflammatory cytokines and chemokines and promoted the cytotoxicity of T(CMV) cells against CMV-infected target cells. Collectively, these observations demonstrate the indispensable role of TBKBP1 for optimal adaptive T cell responses and contribute to our current understanding of CD8^+^ T cell responses to HCMV.

The role of DNA methylation for T cell fate specification has mostly been studied in relation to cytokines and effector molecules (52,73,74). More recently, a number of genome-wide epigenome profiling studies gave first insights into the global changes upon T cell differentiation (23,25,27,28,35,36,56). A seminal comparative study on acute and chronic viral infections demonstrated that naive antigen-specific CD8^+^ T cells are epigenetically regulated and differentiate rapidly into either a memory or exhaustive phenotype upon infection (74). Interestingly, a recent report showed that even during their early phase of differentiation, CD8^+^

T cells responding to acute and chronic infections displayed distinct transcriptional and epigenetic landscapes (75). Among the unique epigenetic signature genes of T(CMV) cells identified in the present study, *KLRD1*, *TBX21* (coding for T-bet), and *ZEB2* have already been reported to regulate functional properties of memory CD8^+^ T cells and were found to be upregulated in chronic infections or exhausted CD8^+^ T cells (14,75–80). Furthermore, we noted that DMRs linked with *S1PR5*, a migratory receptor that is expressed by circulating memory CD8^+^ T cells, but selectively downregulated in tissue-resident memory CD8^+^ T cells (81–83), were demethylated in T(CMV) cells. A recent report on murine tissue-resident memory T cells demonstrated that *S1pr5* induction is directly controlled by T-bet and Zeb2 (81). Additionally, CD8^+^ T cells from chronic infection models of LCMV exhibit a higher frequency of T-bet^+^ cells and higher T-bet expression levels than those from acute infection models (75). These reports further strengthen the findings from our DNA methylation dataset and suggest that in T(CMV) cells, the selectively demethylated genes *TBX21*, *ZEB2*, and *S1PR5* may jointly coordinate the effector function of antigen-specific CD8^+^ T cells.

Moreover, we identified a DMR associated with *TBKBP1* to be strongly demethylated in T(CMV) cells. TBKBP1 was initially reported to be engaged in the TNFα/NF-κB pathway to elicit innate immune responses (58). The antiviral response of the innate immune system relies on various inducible transcription factors such as NF-κB and IRFs, which account for the production of IFNs and pro-inflammatory cytokines (84–86). Studies showed that IκB kinase-I (IKKε), TNFR-associated factor (TRAF)-dependent NF-κB activator, and TBK1 trigger the phosphorylation of IRF3 and IRF7, which are responsible for transactivating type I and type II IFNs (61,62). A recent report demonstrated that TBK1 can be recruited by TBKBP1 to interact with protein kinase C-theta (PKCθ), thereby enabling TBK1 phosphorylation and activation (59). This finding is in accordance with our own observation on TBKBP1-overexpressing CD8^+^ T cells, which upon restimulation showed an enhanced TBK1 phosphorylation that could potentially support downstream signalling pathways. Moreover, various reports showed that phosphorylation of multiple serine residues within regulatory regions of IRF3 and IRF7 orchestrates TBK1 and IκB-mediated signalling pathways (87,88), that IRF7 can promote the expression of IFN-γ-responsive genes through the initiation and stabilization of RNA polymerase II recruitment in the presence of TBK1 and/or IKKε (87), and that IRF3 affects granzyme B expression and maintenance of memory T cell function in response to viral infection (89). These findings are in line with data from the present study demonstrating that ectopic expression of TBKBP1 enhances the cytotoxic activity of CD8^+^ T cells through the upregulation of several pro-inflammatory mediators such as IFN-γ, granzyme A, granzyme B, and perforin. Collectively, our data indicate that TBKBP1 promotes the activation of TBK1 and further downstream pathways for optimal effector function of CMV-specific CD8^+^ T cells.

In conclusion, our findings provide evidence for the potential utility of TBKBP1 as a therapeutic target for the modulation of CD8^+^ T cell-mediated immune responses. Clinical relevance of our findings lies in the fact that ectopic expression of TBKBP1 enhanced the cytotoxicity of T(CMV) cells, suggesting that TBKBP1 could be exploited as a potential therapeutic target for the improvement of adoptive T cell therapy in CMV-infected individuals.

## DATA AVAILABILITY

Sequencing data and methylation levels reported in this paper were uploaded to GEO under accession number GSE245832. Identified DMRs are listed in Supplementary Table 1.

## SUPPLEMENTARY DATA

Supplementary Data are available at NAR online.

## AUTHOR CONTRIBUTIONS

Zheng Yu and Varun Sasidharan Nair: Formal analysis, Investigation, Methodology, Validation, Software, Visualization, Writing-original draft. Agnes Bonifacius: Formal analysis, Visualization, Writing-review & editing. Fawad Khan, Thalea Buchta, Michael Beckstette, Jana Niemz, Philipp Hilgendorf, Beate Pietzsch, Philip Mausberg: Methodology, Validation. Andreas Keller, Christine Falk, Dirk Busch, Melanie M. Brinkmann, Kilian Schober, Luka Cicin-Sain, Fabian Müller: Methodology, Validation, Data curation, Visualization, Resources. Britta Eiz-Vesper: Conceptualization, Resources, Formal analysis, Funding acquisition, Methodology. Stefan Floess: Data curation, Formal analysis, Methodology, Validation, Visualization, Writing-review & editing. Jochen Huehn: Conceptualization, Formal analysis, Funding acquisition, Investigation, Methodology, Supervision, Validation, Writing-review & editing.

## ACKNOWLEDGEMENTS

We thank Lothar Gröbe for carrying out flow cytometry–based cell sorting. We acknowledge the Genome Analytics Facility at the Helmholtz Centre for Infection Research. We appreciate Kerstin Beushausen and Jana Keil from Hannover Medical School for cytokine measurements. We also acknowledge Jana Reinking and Lothar Jänsch for their assistance with protein quantification and CD3/CD28 cross-linking experiments.

## FUNDING

This work was supported by the Life Science Foundation (stipend to ZY), the Friends of the HZI foundation (stipend to VS), the Helmholtz Association (W2/W3-090 to MMB), the German Research Foundation (DFG; as part of the Research Unit 2830 “Advanced Concepts in Cellular Immune Control of Cytomegalovirus”, grant no 431451204 to BEV), and the German Federal Ministry of Education and Science (BMBF; project 01KI2013 to KS and PH).

## CONFLICT OF INTEREST

There are no competing interests declared by the authors.

## Notes

https://www.ncbi.nlm.nih.gov/geo/query/acc.cgi?acc=GSE245832

### Competing Interest Statement

The authors have declared no competing interest.

