## Supplementary Figures for "DNA methylation profiling identifies TBKBP1 as potent amplifier of cytotoxic activity in CMV-specific human CD8^+^ T cells"

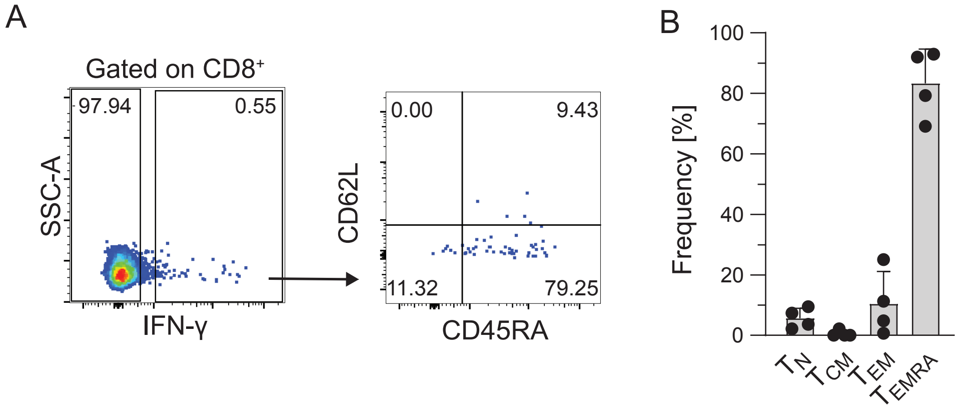


**Supplementary Figure 1: Phenotypic characterisation of T(CMV) cells.** PBMCs from healthy CMV-seropositive donors were pre-enriched for $CD8^{+}$ T cells and subsequently stimulated with the CMVpp65 overlapping peptide pool. Next, IFN-γ-secreting cells were detected using the IFN-γ Secretion Assay/Detection Kit. The phenotype of IFN-γ-secreting T(CMV) cells was determined using flow cytometry. (A) Representative flow cytometry plots show the identification of T(CMV) cells (left) and the phenotypic characterisation via CD45RA and CD62L expression (right). Numbers indicate frequencies in gates or quadrants. (B) The bar plot shows the frequencies of $T_N$ ($CD45RA^{+}CD62L^{+}$), $T_{CM}$ ($CD45RA^{-}CD62L^{+}$), $T_{EM}$ ($CD45RA^{-}CD62L^{-}$) and $T_{EMRA}$ cells ($CD45RA^{+}CD62L^{-}$) within T(CMV) cells from 4 donors. Black dots indicate frequencies from individual donors and grey bar mean values with SD.

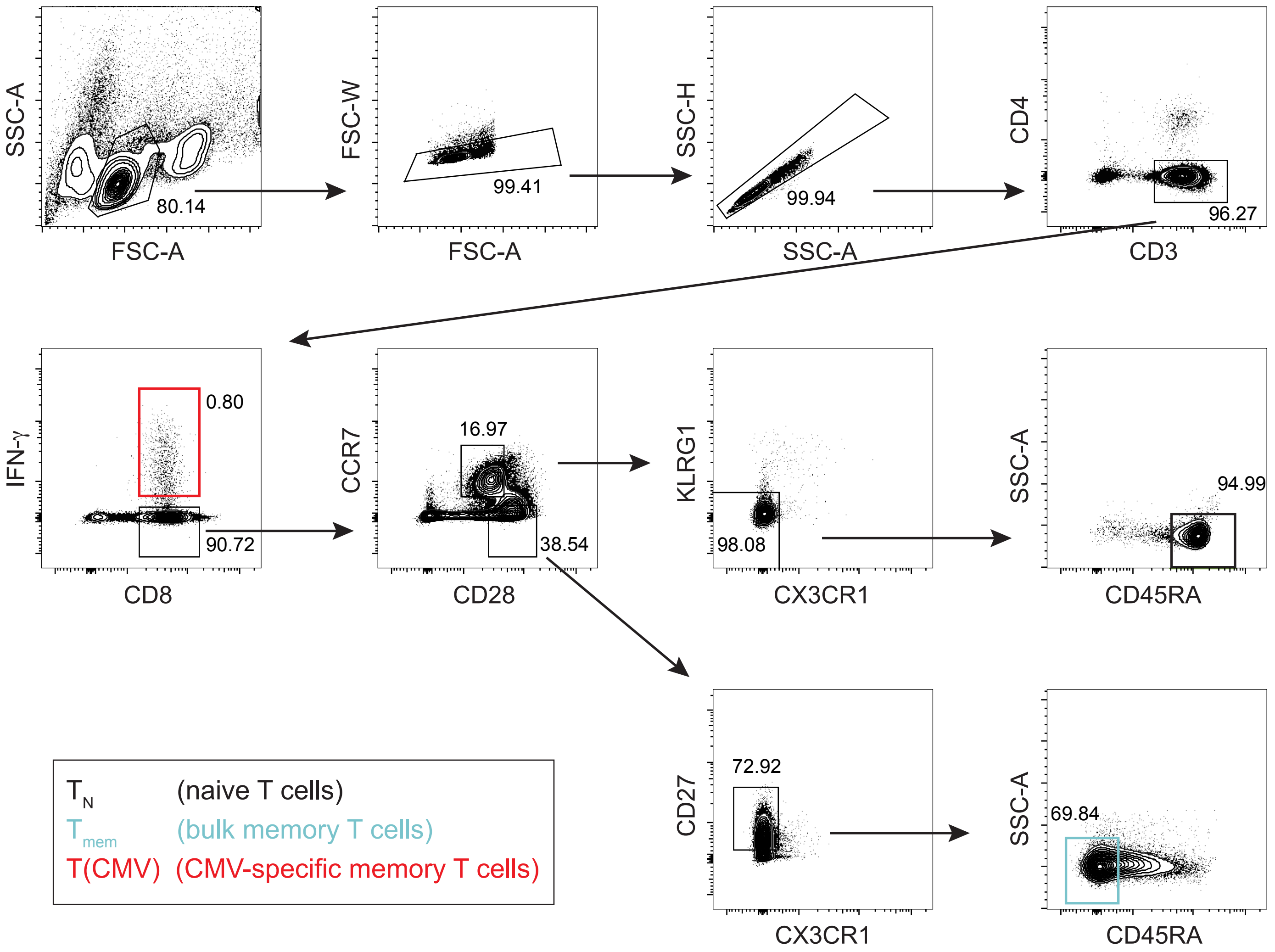


**Supplementary Figure 2: Sorting of CD8$^+$ T cell subsets for WGBS.** PBMCs from healthy CMV-seropositive donors were pre-enriched for CD8$^+$ T cells and stimulated with CMVpp65 overlapping peptide pool to detect IFN-γ-secreting T(CMV) cells using IFN-γ Secretion Assay/Detection Kit. Representative flow cytometric plots show the gating strategy for sorting of $T_N$ (CD3$^+$CD4$^-$CD8$^+$CCR7$^+$CD28$^{int}$KLRG1$^-$CX3CR1$^-$CD45RA$^{high}$), $T_{mem}$ (CD3$^+$CD4$^-$CD8$^+$CCR7$^-$CD28$^{high}$CD27$^+$CD45RA$^-$), and T(CMV) cells (CD3$^+$CD4$^-$CD8$^+$IFN-γ$^+$) from 5 donors.

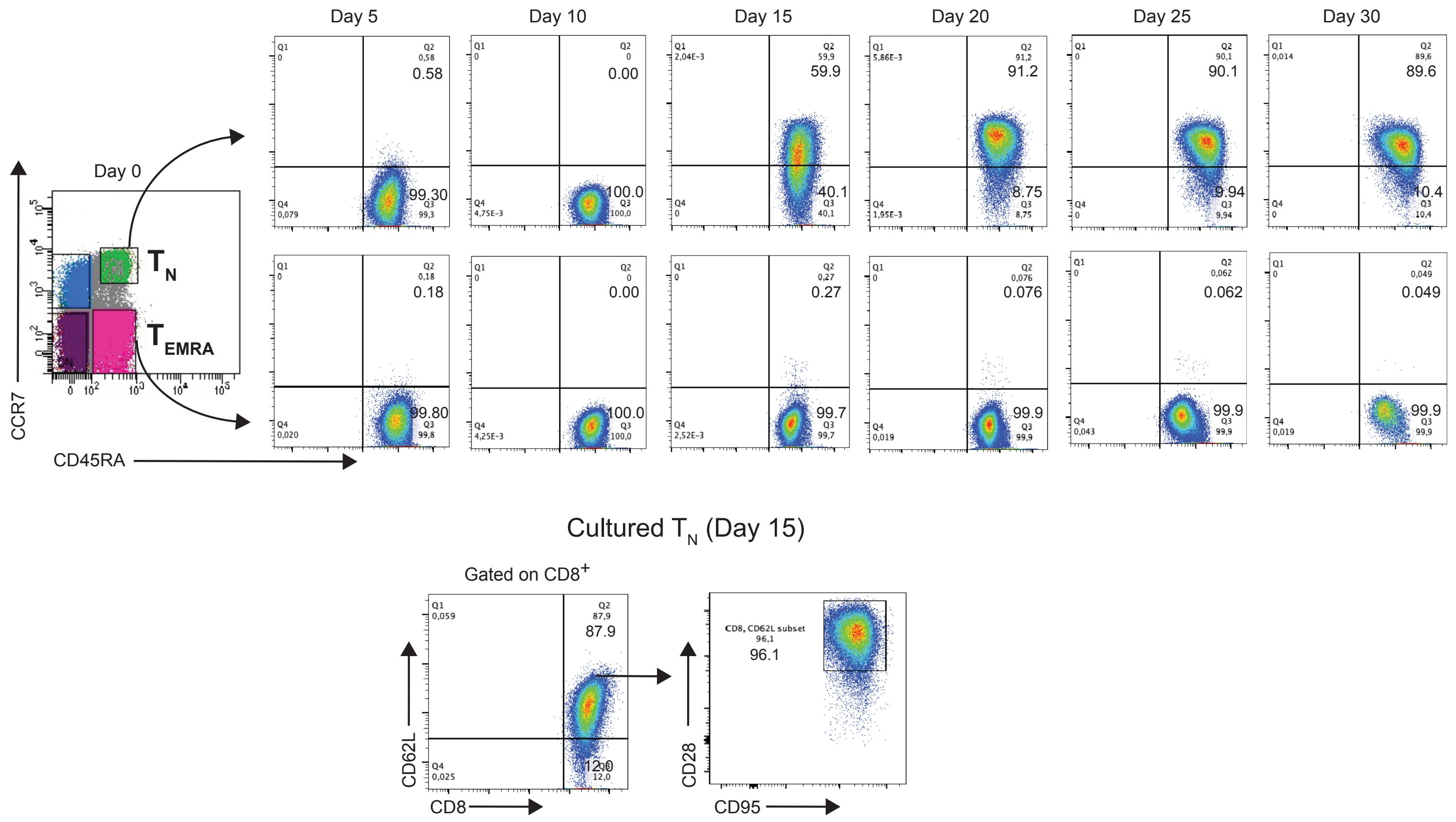


**Supplementary Figure 3: Phenotypic characterisation of CD8$^+$ $T_N$ and $T_{EMRA}$ cells during long-term cultivation.** CD8$^+$ $T_N$ and $T_{EMRA}$ cells were sorted from healthy CMV-seropositive donors and cultured up to 30 days with repetitive restimulations using plate-bound anti-human CD3 and anti-human CD28 antibodies. Every 5 days, cells were harvested, washed and an aliquot was collected to determine the expression of CD8, CCR7, CD45RA, CD62L, CD95, and CD28 by flow cytometry. Representative flow cytometry plots from 5 independent cultures are depicted.

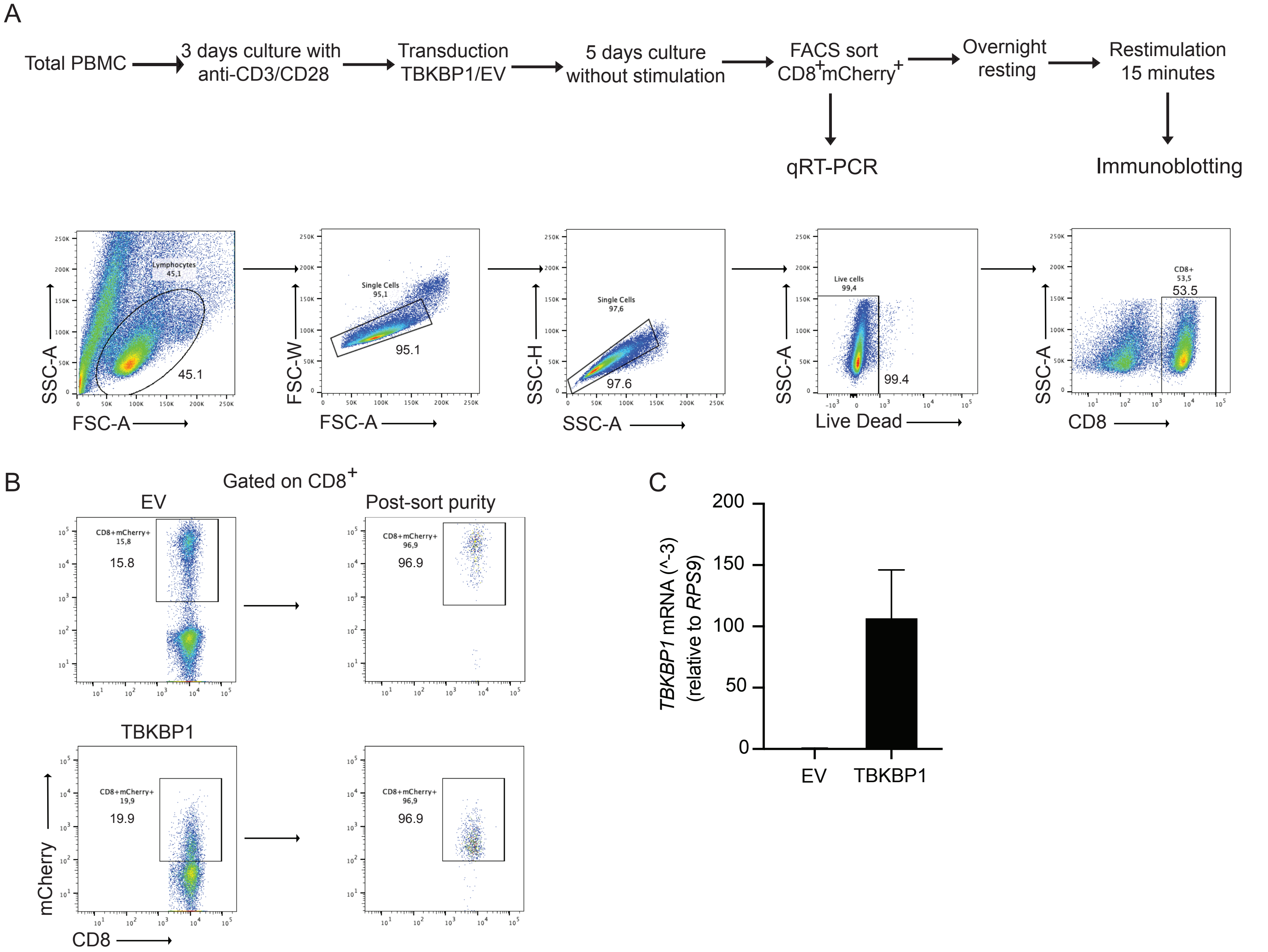


**Supplementary Figure 4: Validation of retroviral TBKBP1-overexpression in $CD8^+$ T cells.** (A) Workflow describes the steps involved in sample processing for immunoblotting experiments. Briefly, after 3 days of activation, PBMCs obtained from CMV-seronegative donors were transduced with pMP71-based vectors. From both empty vector (EV)- and TBKBP1-transduced samples, $CD8^+mCherry^+$ cells were sorted and restimulated with anti-human CD3 and anti-human CD28 for 15 minutes followed by cross linking with goat anti mouse IgG, by keeping unstimulated cells as control. Both unstimulated and restimulated cells from EV and TBKBP1 samples were subjected to immunoblotting to determine the expression levels of TBKBP1, TBK1, pTBK1, and GAPDH. (B) Representative flow cytometry plots show gating strategy for the sorting of $CD8^+mCherry^+$ T cells after overexpressing TBKBP1 and post-sort purity from 4 donors. (C) Bar plots show the mRNA expression of *TBKBP1* in sorted $CD8^+mCherry^+$ T cells from both TBKBP1-overexpressing samples and EV-transduced controls.

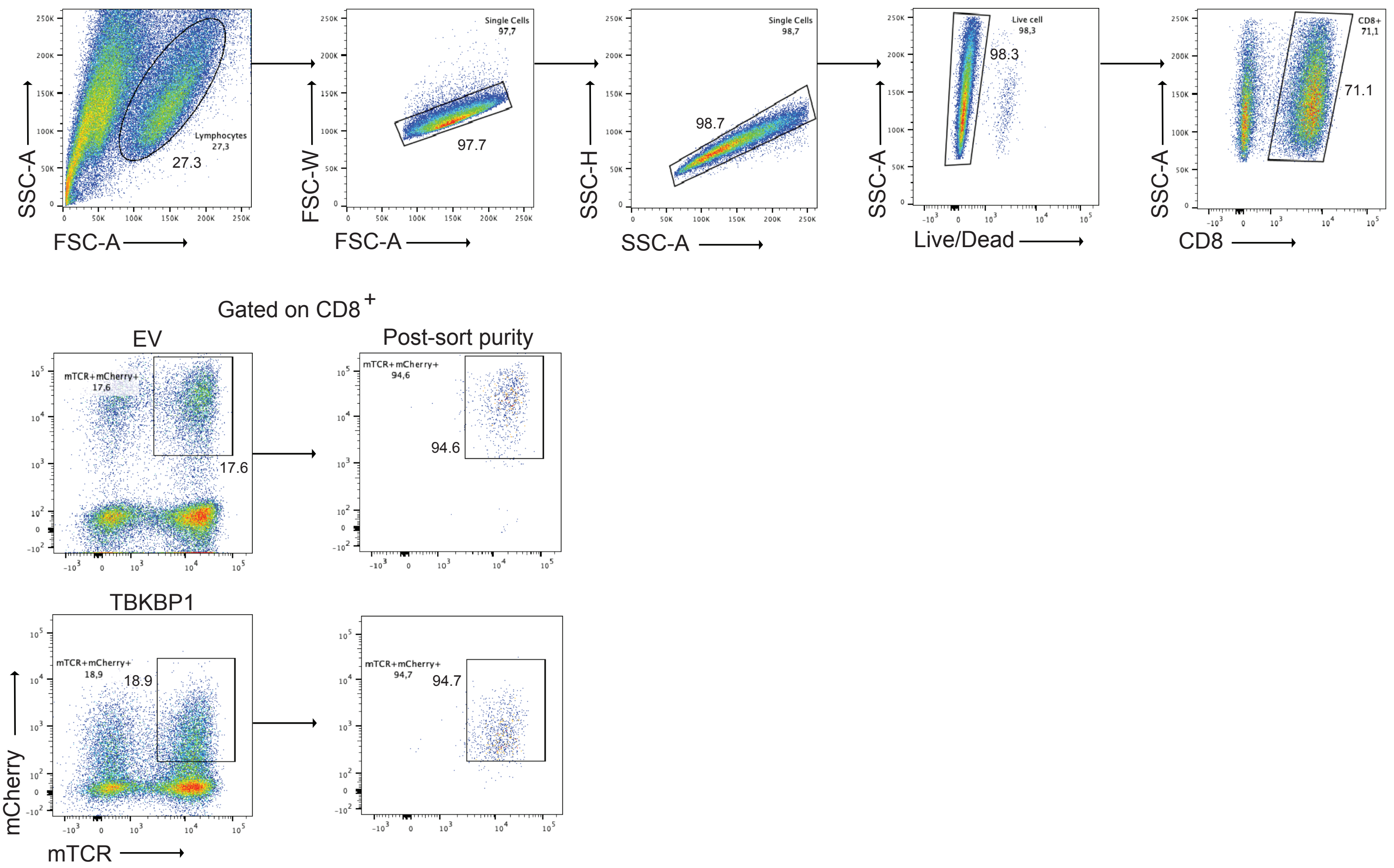


**Supplementary Figure 5: Gating strategy for sorting of TBKBP1-overexpressing CD8$^{+}$ T cells and EV-transduced controls.** PBMCs isolated from healthy CMV-seronegative donors were stimulated with plate-bound anti-human CD3 and anti-human CD28 antibodies and subsequently co-transduced with mTCR and TBKBP1- or EV-mCherry plasmids. Successfully transduced CD8$^{+}$mTCR$^{+}$mCherry$^{+}$ T cells were sorted using flow cytometry. Representative flow cytometry plots from 3 independent donors show the gating strategy for sorting of CD8$^{+}$mTCR$^{+}$mCherry$^{+}$ T cells and post-sort purity.

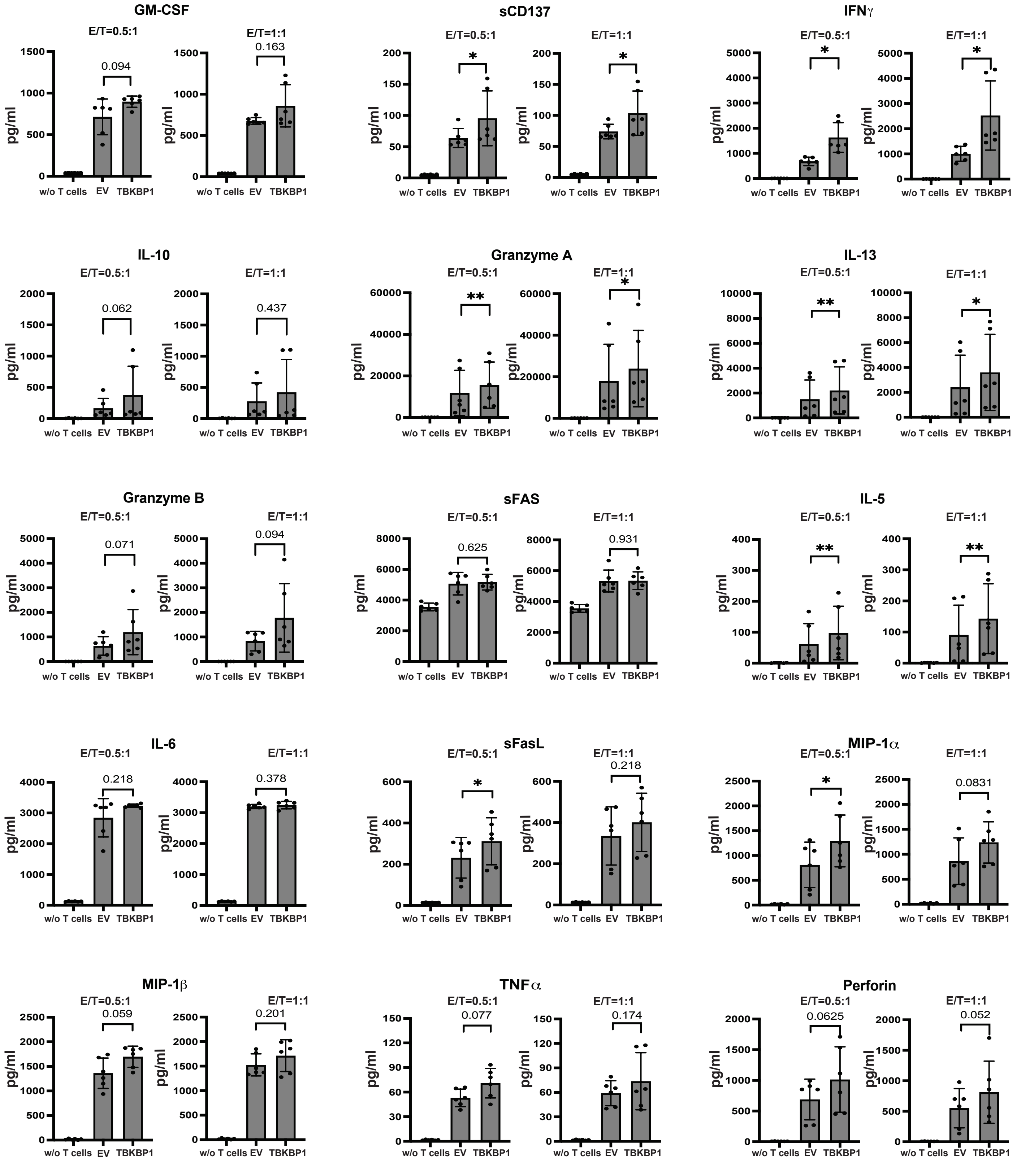


**Supplementary Figure 6: Quantification of cytokines from dynamic virus reduction assay.** PBMCs isolated from healthy CMV-seronegative donors were stimulated with plate-bound anti-human CD3 and anti-human CD28 antibodies and subsequently co-transduced with mTCR and TBKBP1- or EV-mCherry plasmids. Successfully transduced $CD8^+mTCR^+mCherry^+$ T cells were sorted from both TBKBP1-overexpressing samples and EV-transduced controls using flow cytometry and co-cultured with CMV-infected MRC-5 cells followed by the dynamic virus reduction assay. 36 hours after infection, culture supernatants were harvested and cytokine profiles were determined from cultures of CMV-infected MRC-5 cells in the absence of added $CD8^+$ T cells (MRC-5), in the presence of EV-transduced mTCR 5-$2^+$ $CD8^+$ T cells (EV) or in the presence of TBKBP1-overexpressing mTCR 5-$2^+$ $CD8^+$ T cells (TBKBP1) with E:T ratios of 0.5:1 (left) and 1:1 (right) for indicated cytokines. Data from 3 independent experiments with 2 technical replicates each are shown. Black dots indicate frequencies from individual donors and grey bar mean values with SD. For statistical analyses, a paired two-tailed student's *t* test (leaving out "w/o T cell" group) was conducted with *, $p \leq 0.05$ and **, $p \leq 0.01$.
